# Mating status drives fitness trade-offs in exercised female *Drosophila*

**DOI:** 10.1101/2025.06.11.659196

**Authors:** Anne E. Backlund, Meghan Green, Emie Vandiver, Laura K. Reed

## Abstract

Regular physical exercise has been shown to improve physical and psychological well-being through a variety of mechanisms; however, the degree to which different individuals respond to exercise varies depending on sex and genetic factors. *Drosophila* has been used as a model organism to further understand the molecular mechanisms that underlie exercise adaptation. Essential for flies’ ability to adapt to exercise, octopamine is a hormone and neurotransmitter found in invertebrates that is analogous to norepinephrine. Interestingly, octopamine is also crucial for female post mating responses, and no studies to date have explored the interaction between exercise response and reproductive state in females. Here, we investigated the sexual dimorphism in exercise response by exercising male and female flies of multiple *Drosophila* Genetics Reference Panel (DGRP) lines and measuring fitness traits such as climbing ability and starvation resistance. Further, we were interested in how mating status might affect females’ ability to adapt to exercise, and whether the stress of exercise would affect fertility. Our findings show that while male flies are naturally faster climbers than female flies, females tend to be better suited to resist starvation. Additionally, we found that mating status has a significant impact on female flies’ climbing performance and lifespan, and exercise can have negative effects on lifespan and fertility. Surprisingly, we found that exercise has little effect on stored triglycerides, protein levels, or gene expression. DGRP genetic line was also a significant factor that influenced most phenotypes we measured, underscoring the importance of studying multiple genotypes in conjunction with other experimental variables. Results from our study suggest that female flies may experience evolutionary tradeoffs between physical activity, survival, and fertility, and whether the female has mated or not dictates how they respond to physiological stressors such as exercise.

## Introduction

Exercise has numerous health benefits in humans, including increasing bone density and muscle mass [1], reducing blood pressure and inflammation [2–4], and improving insulin sensitivity [5,6]. These benefits are also associated with reduced risk for cardiovascular disease [7,8], metabolic syndrome [9,10], and cancer [11]. Recently, exercise has also been shown to ameliorate cognitive decline and is associated with reduced risk for neurodegenerative diseases such as Alzheimer’s [12–14]. However, the degree to which different individuals respond to exercise varies, with sex and genetics playing large roles in this variation [15–18].

*Drosophila* has been established as a model organism for exercise research due to their genetic similarity to humans and the fact that their short lifespans allow researchers to easily conduct longitudinal studies [19]. Here, we use flies from the *Drosophila* Genetics Reference Panel (DGRP), which are fully sequenced, wild-derived, isogenic inbred lines that allow researchers to study natural genetic variation [20]. DGRP lines have been used in the *Drosophila* exercise field to identify how genetic background influences phenotypes such as activity, climbing ability, metabolic traits, and gene expression [21,22]. One device that is utilized to induce exercise in *Drosophila* is called the Power Tower, which subjects flies to a repetitive dropping motion that knocks the flies to the bottom of the vial, taking advantage of their negative geotaxis [23]. A previous study using the Power Tower found that male flies can improve their endurance, flight performance, climbing speed, and cardiovascular health after three weeks of exercise training on the Power Tower, while females are unable to gain these positive adaptations [24]. Sujkowski et al.’s findings revealed that octopamine (OA) – a hormone and neurotransmitter analogous to norepinephrine in vertebrates – was essential for flies’ ability to adapt to exercise. Unexercised flies fed OA-supplemented food performed similarly to exercised flies, suggesting that increased OA levels can mimic the effects of exercise. Further, their results proposed that differential activity of octopaminergic neurons was the reason why females seem to be “resistant” to exercise [24].

OA has been studied in *Drosophila* and other insects as it is a critical regulator for many processes such as locomotion [25], flight [26], metabolism [27,28], and interestingly, female reproduction [29–31]. This led us to question how exercise and female fertility are linked, as they are both governed by OA. Exercise and reproduction are energetically costly processes, requiring global changes in metabolism and gene expression [29]. Octopamine is a known metabolic regulator, so we hypothesized that octopamine mediates the shift in metabolism required for females to allocate energetic resources to either reproduction or exercise.

The goal of our study was to investigate sex differences in exercise response in *Drosophila,* and specifically how exercise can affect female fertility and fitness traits. Additionally, we wanted to understand how OA may be involved in these processes, as previous research has studied these ideas independently, but there are no studies to date that examine them together. First, we exercised male and female flies from six different DGRP lines and explored how OA-supplemented food might affect their climbing performance and starvation resistance. Next, we investigated how female flies’ mating status would influence their ability to adapt to exercise by exercising age-matched virgin and mated females from three different DGRP lines and measuring their climbing, lifespan, triglyceride and protein levels, and gene expression. We also explored how exercise would impact mated females’ fertility during and after exercise.

## Materials and methods

### Fly Stocks and Maintenance

The Drosophila Genetic Reference Panel stocks were obtained from the Bloomington Stock Center (genetic lines 301 (stock ID #25175), 324 (#25182), 399 (#25192), 437 (#25194), 900 (#28261), and 907 (#28262)). These lines are all naturally *Wolbachia* negative to control for any interaction effects that may occur between this endosymbiont and the host [32]. For Experiment 1, 2-3 different genetic lines were tested together in blocks over time to control for any batch effects. For Experiment 2, all flies from the three genetic lines used (324, 399, and 900) were tested at once. For both experiments, groups of 50 first instar larvae were transferred onto standard cornmeal-molasses lab food (by weight 5.28% cornmeal, 1.05% yeast, 0.56% agar, 87.03% water, 4.37% molasses, 1.15% Tegosept, 0.55% Propionic acid) and were raised in incubators at 25 °C, 50% relative humidity, on a 12h:12h light:dark cycle. For Experiment 1 (Fig 1A), experimental flies were collected four days post-eclosion (female flies presumed to be mated) under brief CO_2_ and placed on food with a higher agar ratio (10% sucrose, 10% yeast, and 2% agar in water) which provides flies with a cushion when exercising on the Power Tower. A subset of each exercise treatment group was fed OA-supplemented food (5 mg/L, see Octopamine Dosage Experiment methods section for more information) for three hours a day (placed on food one hour before exercise and removed one hour after exercise). OA food was only given to the flies for a few hours to simulate the effects of one bout of exercise, as opposed to having consistently high levels of OA, which could be indicative of chronic stress. Cohorts of 12 flies per vial were randomly assigned to treatment groups and began the exercise experiment the following day (five days post-eclosion). There were six biological replicates per treatment group per genetic line (n=72 flies per treatment). For Experiment 2 (Fig 1B), virgin female flies were collected within eight hours of eclosion under brief CO_2_, placed in food vials (10% sucrose, 10% yeast, and 2% agar in water), and kept in the incubator until the experiment began. Mated female flies were kept with males for four days to ensure that they had mated and then were collected under brief CO_2_ and placed in food vials. Used vials of food were kept in the incubator for a week after the flies were transferred to new food to validate that the virgin females produced no offspring and the mated females did produce offspring. Cohorts of approximately 15 flies per vial were randomly assigned to treatment groups and all flies began the exercise experiment at five days post-eclosion. There were nine biological replicates per treatment group per genetic line (n=135 flies per treatment).

**Fig 1.**
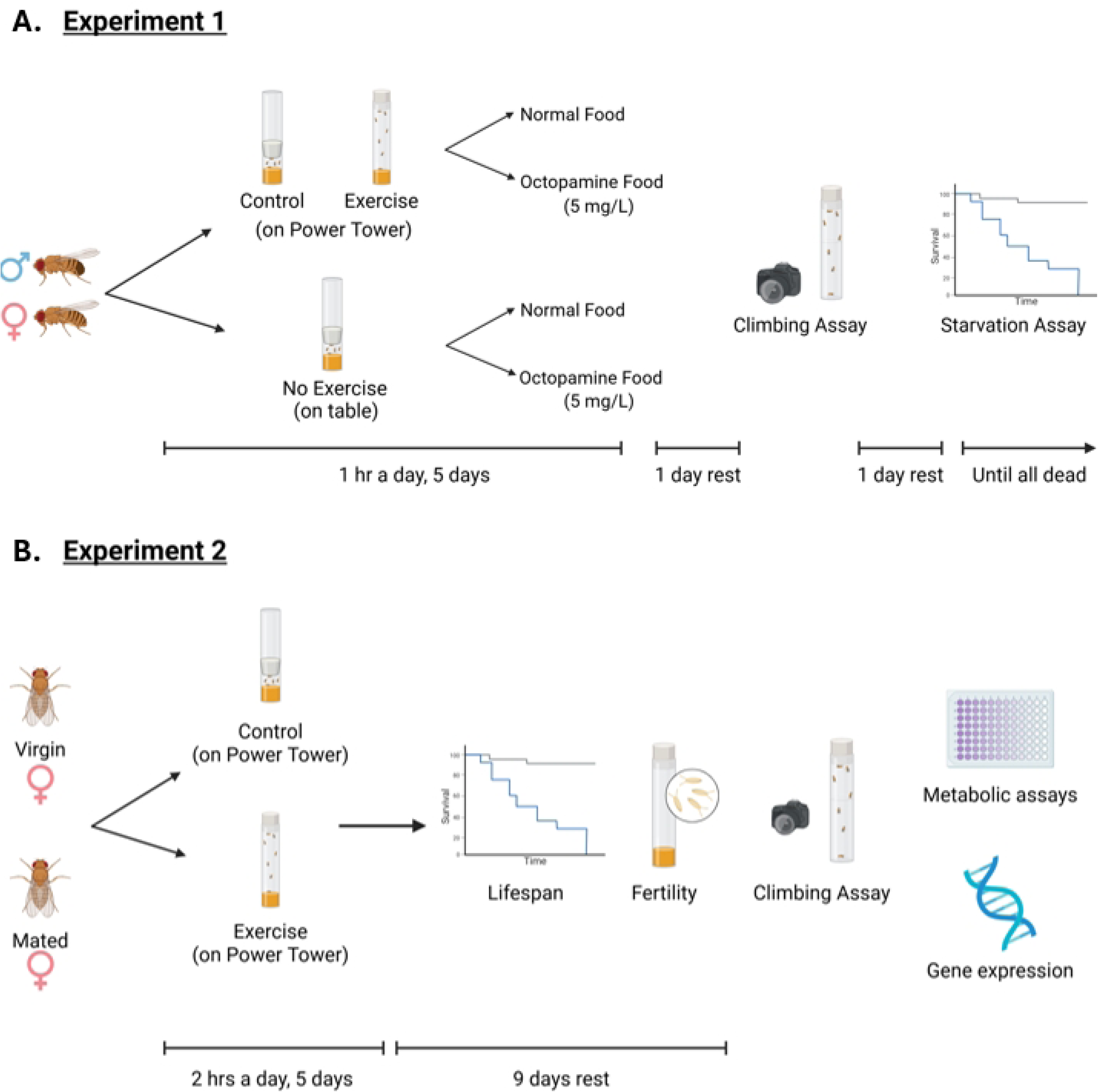
Experimental design diagram. Created in BioRender. **Backlund, A. (2025) https://BioRender.com/8h3iqbq** (A) Experimental design for Experiment 1. (B) Experimental design for Experiment 2.

### Exercise Training

For Experiment 1, flies assigned to the Exercise treatment group were placed on the Power Tower to exercise for one hour a day for five consecutive days, with the Power Tower dropping the flies every 15 seconds (Fig 1A). Control flies were also placed on the Power Tower with their foam stopper pushed down to ½ cm above the level of the food to limit the flies’ ability to move. An additional control group that we will refer to as the “No Exercise” group also had their foam stopper pushed down and was placed on a bench near the Power Tower where they remained stationary for the entire exercise period. After exercise treatment, flies were returned to the incubator for the rest of the day. Flies were transferred onto fresh food every two days and the number of dead flies was recorded. The climbing assay (described below) was performed two days after the cessation of exercise, then flies were promptly returned to food vials and given one more day of rest before beginning the starvation assay.

For Experiment 2, exercised flies were placed on the Power Tower to exercise for two hours a day for five consecutive days (Fig 1B). Control flies were also placed on the Power Tower with their foam stopper pushed down to approximately ½ cm above the level of the food. The “No Exercise” group was not used in Experiment 2 as there were no statistically significant differences between the Control and No Exercise groups in Experiment 1. After exercise treatment, flies were returned to the incubator. After five days of exercise, flies were kept alive for another nine days to continue tracking lifespan and fertility. At 18 days old, the flies performed the climbing assay (described below) and then were flash-frozen in liquid nitrogen and stored at −80°C for later metabolic assays and qRT-PCR.

### Climbing Assay

Climbing ability was measured via a negative geotaxis assay (adapted from [33]). A digital camera was placed at a fixed location six inches away from flies in an empty polystyrene fly vial (9.5 cm long). A piece of paper with a 1 cm x 1 cm grid was placed behind the vial to provide a scale for analysis. The vial was firmly tapped on the table three times, knocking the flies to the bottom of the vial, and then a photograph was taken of the flies four seconds later using a timer on the camera. Four seconds was chosen as an appropriate length of time to be able to see variation in climbing height without all the flies reaching the top of the vial. Each vial of flies was assayed three consecutive times, and the images were analyzed using Fiji (ImageJ) software to measure how far each fly climbed from the bottom of the vial [34]. Average climbing height was calculated for each picture using a macro in Fiji which allowed the user to log flies’ positions in the images and calculate their climbing heights from the bottom of the vial.

### Starvation Assay

To quantify flies’ fitness and ability to resist starvation, flies were placed in vials containing 1% w/v agar and water. The number of dead flies was recorded every 24 hours until all flies were dead. Flies that escaped or were accidentally killed were censored from the survival analysis.

### Octopamine Dosage Experiment

We performed a small experiment testing different concentrations of octopamine to determine if a higher dose than what was originally used (5 mg/L) would be sufficient to increase climbing heights even in unexercised flies. Octopamine hydrochloride (Sigma Aldrich) was dissolved in 10 mL water to produce stock solutions of varying concentrations (5, 50, 500, 5000 mg/L) which was then added to the fly food recipe (10% sucrose, 10% yeast, and 2% agar in 770 mL of water). Normal food with an equal volume of water was used as a control. Male and virgin female flies from DGRP lines 324 and 900 were fed these diets continuously for three weeks (starting from day one post-eclosion) and then a climbing assay was performed as described above.

### Lifespan and Fertility

For Experiment 2, flies were transferred onto fresh food twice a week and the number of dead flies was recorded each day. Flies that escaped or were accidentally killed were censored from the survival analysis. To measure fertility, used vials from mated females were kept in the incubator and the number of pupae in each vial was counted after seven days. The number of pupae was divided by the number of females in the vial and the number of days the females were in the food vial to obtain a normalized value of pupae per female per day.

### Metabolic Assays

After flash-freezing, flies from each treatment group were randomly sorted into groups of 10 for metabolic assays and qRT-PCR. Triglyceride concentrations were measured by absorbance using the Sigma Serum Triglyceride Determination Kit (TR0100) as described in Mendez et al. 2016 [21], with three biological replicates per treatment (n=10 flies per replicate). Protein concentrations were measured using the Bradford method to normalize the triglyceride results to the flies’ approximate size [35].

### qRT-PCR

RNA was extracted from flies (three biological replicates per treatment, n=10 flies per replicate) using the Omega Bio-tek E.Z.N.A.® Total RNA Kit I according to the company protocol. cDNA was synthesized using iScript Reverse Transcription Supermix for RT-qPCR according to the kit protocol. qRT-PCR was performed on these cDNA samples (diluted to 20 ng/µL) using BioRad SYBR green fluorescent dye. The reaction was run under the following conditions: enzyme activation 95 °C for 30 seconds, 40 cycles of denaturation 95°C for 10 seconds and annealing/extension 60°C for 30 seconds, melt curve 65-95°C in 0.5°C increments for 5 seconds each step. Expression of the housekeeping gene *Rp49* was used to calculate ΔCT, and ΔΔCT was calculated by subtracting ΔCT of a pooled sample from ΔCT of each individual sample. The following primers were used: *AkhR*-f: 5’-GGACTCTACAACATTCGC-3’, *AkhR*-r: 5’-CCTCTTCCATTCAGCAGC-3’, *Dilp2*-f: 5’-CTGAGTATGGTGTGCGAGGA-3’, *Dilp2*-r: 5’-CAGCCAGGGAATTGAGTACAC-3’, *Tbh*-f: 5’- ATCCGTACGTTCGACTGGAG-3’, *Tbh*-r: 5’-TCGACATCTTGATGCGAAAG-3’, *Tdc1*-f: 5’- TTGAGGTTCGCAACGATGTTC-3’, *Tdc1*-r: 5’-AAGCACTTTATCTGGGTCCAAGC-3’, *Rp49*-f: 5’- GGTGAGTGCCAACGAGGATT-3’, *Rp49*-r: 5’-CTTGCGCTTCTTGGAGGAGA-3’.

### Statistical Analysis

Data was analyzed using JMP Pro 17. Triglyceride and fertility data were log_10_ transformed for normality prior to analysis, all other phenotypes did not require transformation. To check for the contributions of the experimental variables (e.g., exercise treatment, genetic line, sex, mating status, and their interactions) on the measured phenotypes of climbing height, fertility, triglyceride concentration, protein concentration, and gene expression, we employed a general linear model. We also nested replicate effects within the other experimental factors to account for our experimental design. *Post hoc* tests for pairwise contrasts were performed using a two-tailed Student’s t-test. All p-values were considered significant at p<0.05. All data for presented analyses are available in Supplemental Files 1-9.

Starvation assay and lifespan data were analyzed using Survival Analysis on JMP and p-values were calculated with a Log-Rank test between groups. Flies that escaped or were accidentally killed were censored from the survival analysis. For the octopamine dosage experiment, individual fly climbing heights were analyzed instead of the average climbing height per vial, then these values were converted to binary values depending on if the fly climbed above or below the median climbing height of 2.5 cm. As we did not see a strong relationship between octopamine dosage and climbing height alone, we also analyzed the dosages by level: “low” (0 and 5 mg/L) or “high” (50, 500, and 5000 mg/L). These data were analyzed using a nominal logistic fit model.

## Results

### Experiment 1

To investigate sex differences in exercise response and how OA could be involved, we first exercised male and female flies from six different genetic lines for one hour a day for five days. Results of our overall effect tests show that genetic line (p<0.0001) and sex (p<0.0001) have the largest impacts on climbing height. We also found that in every DGRP line we studied, males climb significantly higher than females overall (Fig 2A). However, when we analyzed how each sex responded to exercise, we discovered that males – although they seem to be naturally faster climbers than females – showed no significant changes in climbing height post-exercise. On the other hand, exercised female flies showed increased climbing heights compared to the Control and No Exercise groups (Fig 2B). Our findings were contrary to those found in Sujkowski et al., which found that females are “resistant” to exercise, whereas males are able to obtain positive adaptations [24]. This difference could be due to the different exercise training programs used, as the other study exercised flies for three weeks with increasing duration each week, and our study only exercised the flies for one hour a day for five days. Additionally, flies’ climbing scores tend to vary over their lifetimes, so the age of the fly in relation to the exercise experiment could impact climbing assay results. Our lab has found that exercise studies using *Drosophila* can be difficult to replicate due to differences in exercise machines, training durations, genetic background, and other environmental factors. We found that exercise treatment group had a significant effect on climbing height overall (p=0.0009), with exercised flies climbing higher than both the Control (on device) and No Exercise (off device) groups (Fig 2C). As there was no significant difference between the No Exercise and Control groups, we opted to not use the No Exercise group in our second experiment.

**Fig 2.**
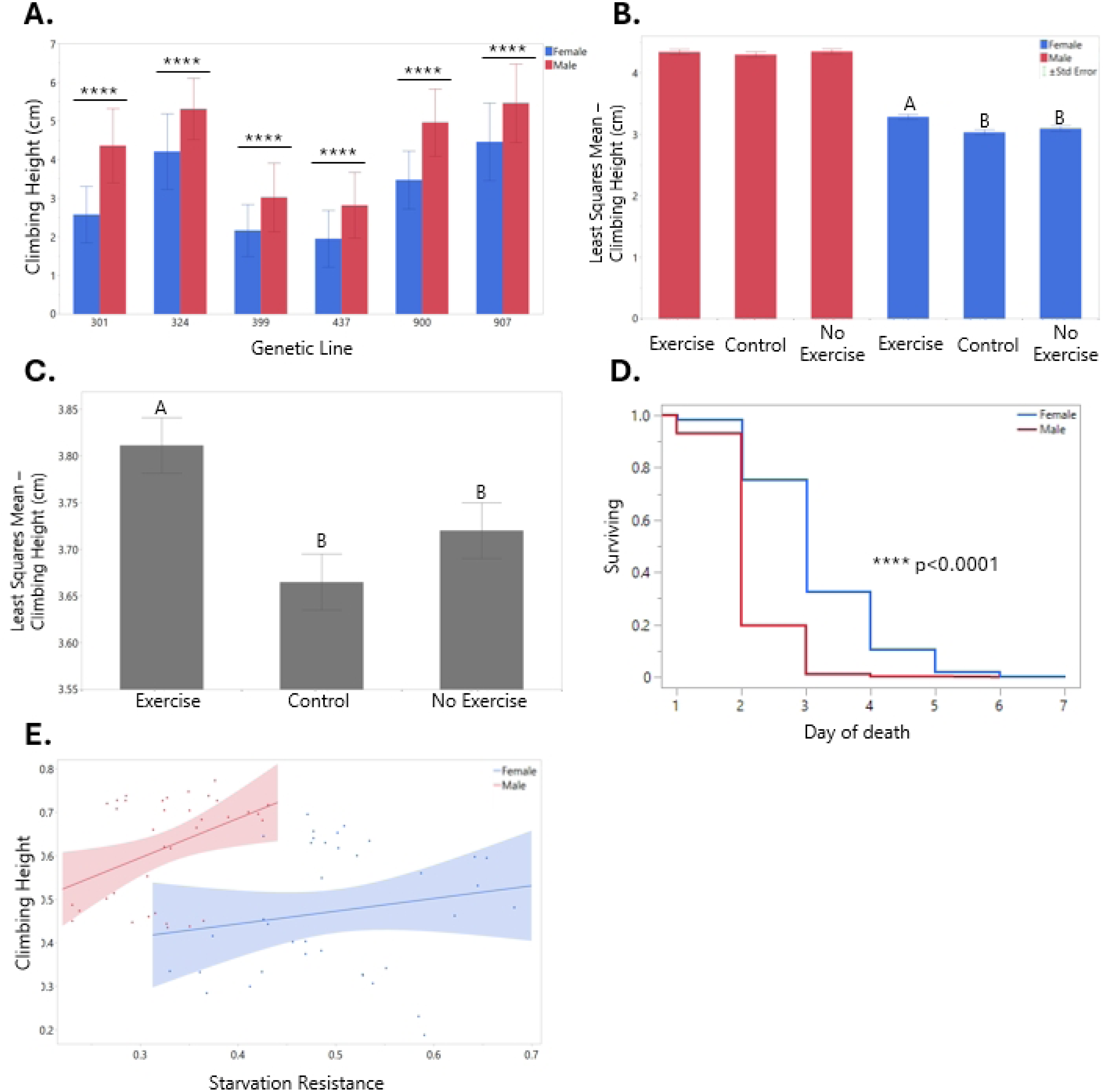
Sex and genotype drive variation in climbing performance and starvation resistance. (A) Climbing assay results for females (blue) and males (red) per genetic line. Error bars are one standard deviation. For each t-test between males and females, p<0.0001. (B) Least squares mean climbing assay results for exercise treatment groups by sex. Error bars are standard error. There were no significant differences between exercise treatment groups for male flies (red). For females (blue), different letters above bars indicate that the values are significantly different (p<0.05), letters that are the same indicate that the values are not significantly different from each other. (C) Least squares mean climbing assay results for the exercise treatment groups. Error bars are standard error. Different letters above bars indicate that the values are significantly different (p<0.05), letters that are the same indicate that the values are not significantly different from each other. (D) Starvation assay results for females (blue) and males (red). (E) Correlation between starvation resistance and climbing height for females (in blue, R=0.172, ns) and males (in red, R=0.405, p=0.0142).

We also hypothesized that flies fed food supplemented with 5 mg/L OA would have greater climbing performance than those fed normal food; however, diet was not a significant main effect. There were many interaction effects, including line-by-sex, line-by-exercise treatment, and line-by-sex-by-diet (Table 1), which illustrates the complexity of genetic and environmental interactions and how they can affect flies’ response to exercise.

**Table 1.** Effect test results for Experiment 1 climbing assay data.

|  |  |
| --- | --- |
| Line | **p<0.0001** |
| Sex | **p<0.0001** |
| Exercise | **p=0.0009** |
| Diet | ns |
| Line * Sex | **p<0.0001** |
| Line * Exercise | **p<0.0001** |
| Sex * Exercise | ns |
| Line * Sex * Exercise | **p<0.0001** |
| Line * Diet | **p=0.0004** |
| Sex * Diet | **p=0.0257** |
| Line * Sex * Diet | **p=0.0005** |
| Exercise * Diet | ns |
| Line * Exercise * Diet | **p<0.0001** |
| Sex * Exercise * Diet | ns |
| Line * Sex * Exercise * Diet | **p=0.0195** |
| Biological replicate [Line, Sex,<br>Exercise, Diet] | **p<0.0001** |
| ID [Line, Sex, Exercise, Diet,<br>Biological replicate] | **p<0.0001** |

Bolded p-values are considered significant (p<0.05).

**Table 2.** Effect test results for Experiment 2 climbing assay data.

| Line | Exercise | Mating Status | Line * Exercise | Line * Mating Status | Exercise * Mating Status | Line * Exercise * Mating Status |
| --- | --- | --- | --- | --- | --- | --- |
| **p<0.0001** | **p<0.0001** | **p<0.0001** | **p<0.0001** | **p<0.0001** | **p<0.0001** | **p<0.0001** |

Bolded p-values are considered significant ( $p<0.05$ ).

After the climbing assay, flies were subjected to a starvation assay to test their overall fitness and lifespan. We found that females exhibit enhanced starvation resistance compared to males (Fig 2D), which makes sense as females are generally larger and store more fat than males [36,37]. Genetic line was also a significant indicator of starvation resistance (p<0.0001), while diet (octopamine feeding) was not a significant main effect. The No Exercise group survived slightly longer than the Exercise and Control groups (p=0.0224). This suggests that the Exercise and Control groups experience stress by being placed on the Power Tower that may leave them more vulnerable to other stressors, such as starvation. Additionally, these flies may have used up their stores of carbohydrates and fats as fuel for exercise, with not enough left over to resist starvation.

The main findings from Experiment 1 were that sex and genetic line strongly impact phenotypes such as climbing ability and starvation resistance in *Drosophila*. We observed that females did not climb as high as males in every genetic line we studied, but females were able to resist starvation for longer than males, leading us to wonder whether there is a correlation between these phenotypes. By plotting starvation and climbing height on the same graph, we were able to see that both males and females have a slight positive correlation between the two variables, but that they clustered in different areas (Fig 2E). This points to a possible evolutionary trade-off that occurs in males and females: males are faster climbers but do not live as long, while females are slower climbers but live longer.

To determine if a higher concentration or longer duration of feeding OA would produce different results, we fed male and female flies different dosages of OA (0, 5, 50, 500, or 5000 mg/L) continuously for three weeks and then performed a climbing assay. We did not observe a clear effect of OA dosage unless we analyzed them by dosage level – low (0 or 5mg/L) or high (50, 500, or 5000 mg/L) – and looked at if the flies could climb higher than the median climbing height (2.5 cm). Using this method of analysis, we found that more flies climbed higher than this threshold after being subjected to the high dose compared to the low dose (p=0.0095) (Fig 3). We also found that genetic line and the interaction between line and sex were significant main effects (p=0.0158 and p=0.0002, respectively). As we did not see a robust effect of dosage, we are skeptical about the general effectiveness of using OA feeding as a research strategy to increase climbing height of *Drosophila*.

**Fig 3.**
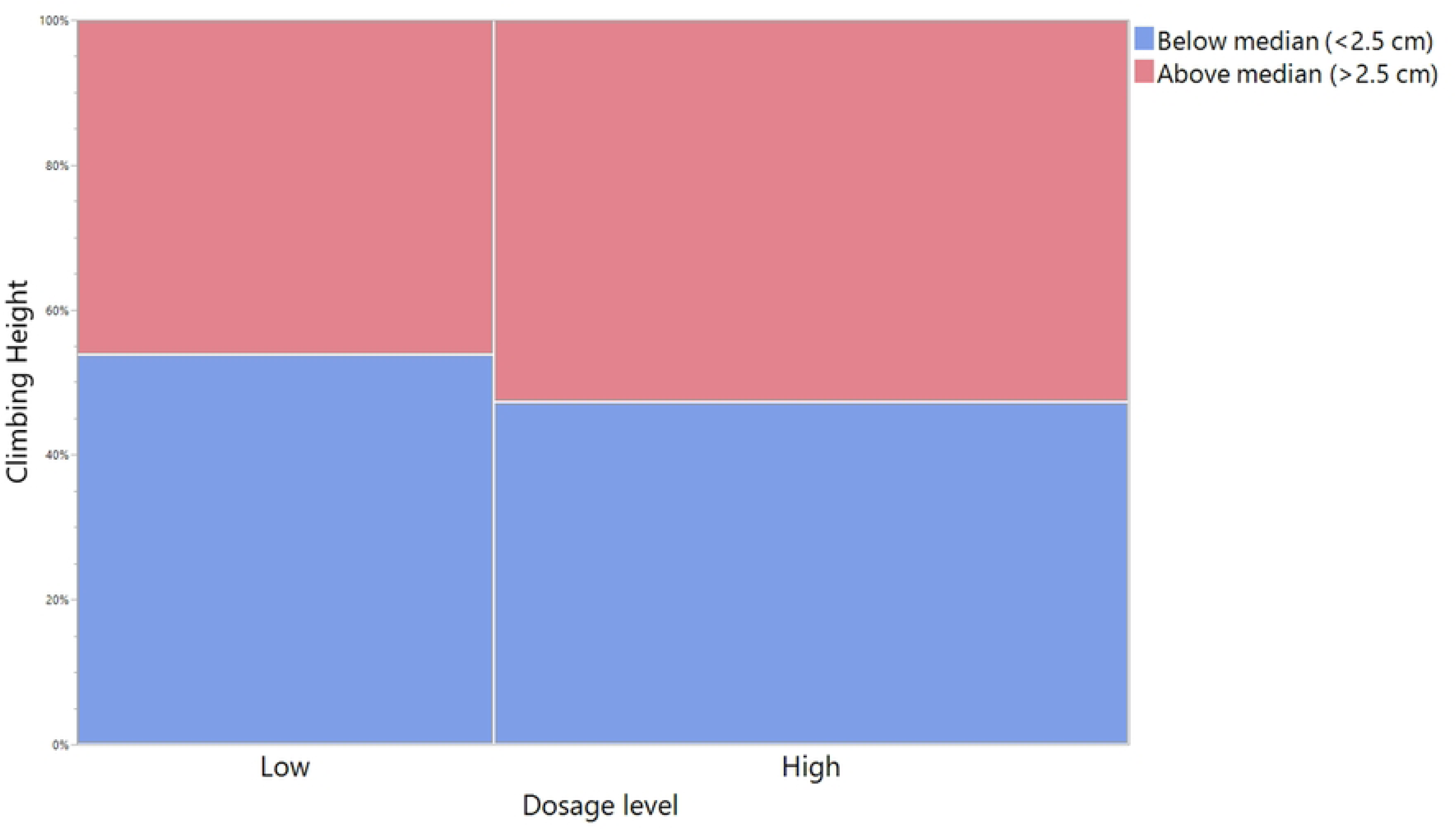
Higher concentrations of octopamine are needed to elicit responses in *Drosophila* climbing performance. OA dosage levels are low (0 and 5 mg/L) and high (50, 500, 5000 mg/L). Climbing height is illustrated as the percentage of flies climbing below the median height of 2.5 cm (blue) or above the median (red).

### Experiment 2

Our findings from Experiment 1 led us to examine whether female *Drosophila* experience a trade-off between survival, exercise adaptations, and/or reproduction. In Experiment 2, we exercised age-matched virgin and mated female flies from three different DGRP lines and measured climbing performance, lifespan, and fertility (for mated females only). Our effect tests for the climbing assay results showed that genetic line, mating status, exercise treatment, and their interactions were significant (Table 3). After nine days of rest, Control flies climbed higher than Exercised flies and virgins climbed higher than mated females. When we looked at these variables in combination, we observed that mated females in the Control group climbed higher than those in the Exercise group, while there was no significant difference between exercise treatment groups for virgin females (Fig 4A). These results suggest that mated females are more susceptible to the stress of exercise. However, the lifespan analysis showed somewhat conflicting results, with both Control and Exercised mated females surviving significantly longer than virgin females (Fig 4B). When analyzing the flies’ lifespans for the length of the experiment, exercise treatment was not significant, but when we only looked at the five days that the flies were exercising, we found that Control flies had significantly greater survival than Exercised flies (Fig 4C). Taken together, this data suggests that virgin female flies may be more physically active than mated female flies, but that this comes at a significant cost to their lifespan. We also hypothesize that mated female flies may be poorer climbers in general because they are allocating more energy to reproduction than physical activity. Finally, we found that exercise is a stressful experience for female flies in the short term, but that there appear to be no long-lasting effects of exercise on lifespan.

**Fig 4.**
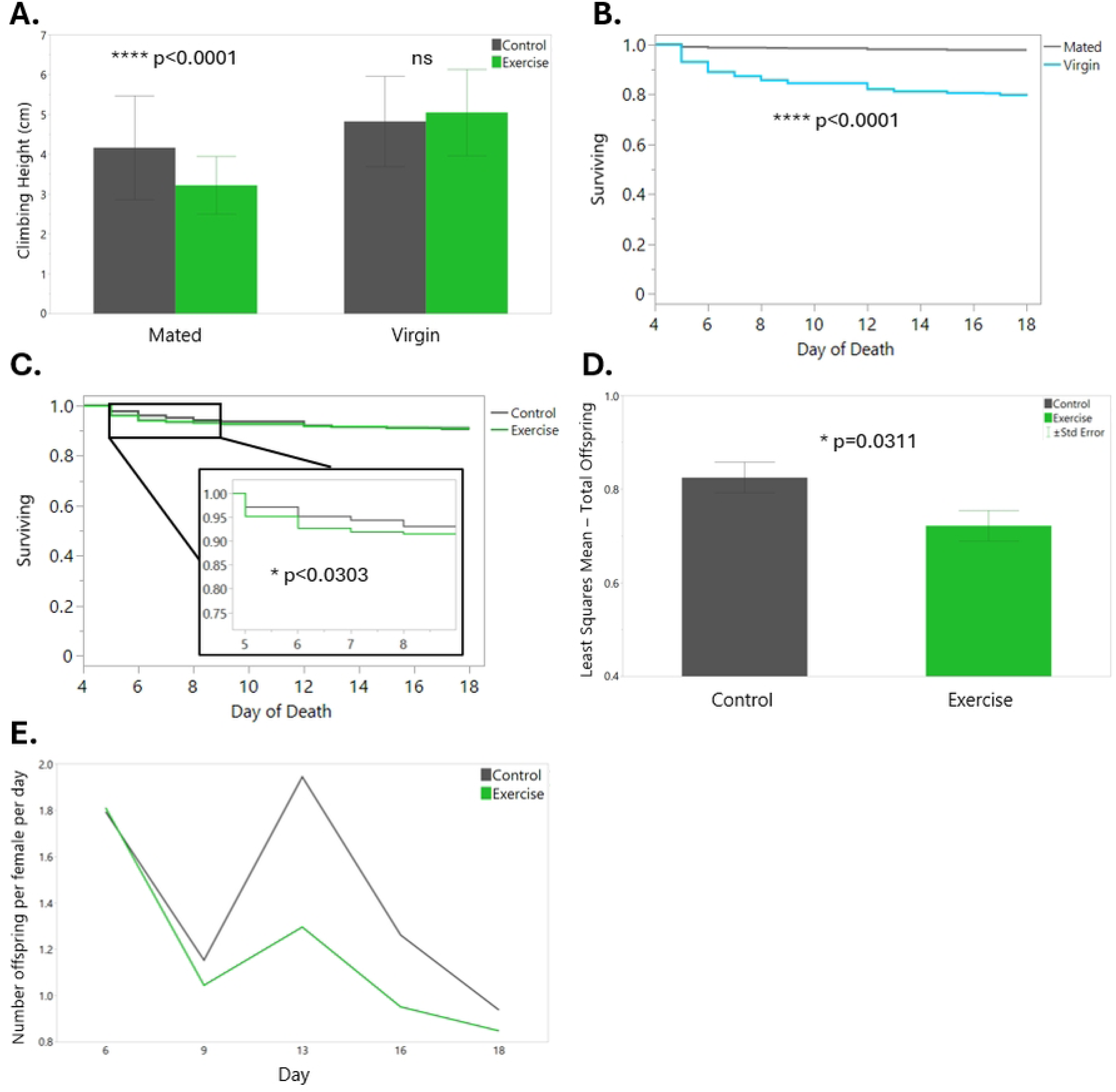
Mating status impacts climbing performance and survival in female flies, and exercise negatively impacts acute survival. Exercise training is also associated with a reduction in fertility. (A) Climbing assay results for Control (gray) and Exercise (green) mated and virgin female flies. Error bars are one standard deviation. (B) Lifespan of mated (gray) and virgin (light blue) female flies. (C) Lifespan of Control (gray) and Exercise (green) female flies with inset showing difference in survival during the 5 days of exercise. (D) Least squares mean total number of offspring for Control (gray) and Exercise (green). Error bars are standard error. (E) Normalized number of offspring per female per day for Control (gray) and Exercise (green) over time.

To further investigate the effects of exercise on fitness of female *Drosophila*, we tracked the number of offspring produced by mated females during and after exercise treatment. Genetic line and exercise treatment were both significant predictors for total offspring (p=0.0014 and p=0.0311, respectively), with Control flies having significantly more offspring than Exercised flies (Fig 4D). This effect lasted more than a week past the cessation of exercise (Fig 4E). Again, these results indicate that exercise is a stressful experience for mated female flies and can decrease their reproductive fitness.

To test the hypothesis that exercised and/or virgin female flies have reduced energy stores, we performed triglyceride and protein assays. There were no significant differences in triglyceride levels between any of the treatment groups, which was a surprising result, as previous studies have found that exercised flies have reduced triglyceride levels [21]. We observed the opposite pattern within DGRP line 900, with Exercised flies having significantly higher triglyceride levels than Control (Fig 5A, p=0.0398). This could be because the flies had nine days of rest before we measured their triglycerides, and during this time the Exercised flies may have consumed more food than the Control flies. Genetic line also had a significant impact on triglyceride concentrations (p=0.0038). Exercise treatment also did not appear to be a significant predictor of protein levels, although the interaction between line and exercise treatment was significant (p=0.0184). This is also contrary to the findings of the previously mentioned paper, which showed that Exercised flies had increased protein levels compared to Control [21]. Mating status did have a significant effect on protein levels, as our results showed that virgin female flies have higher protein concentrations than mated female flies (Fig 5B, p<0.0001). Future studies should investigate other metabolic markers in *Drosophila* post-exercise to learn more about how flies respond.

**Fig 5.**
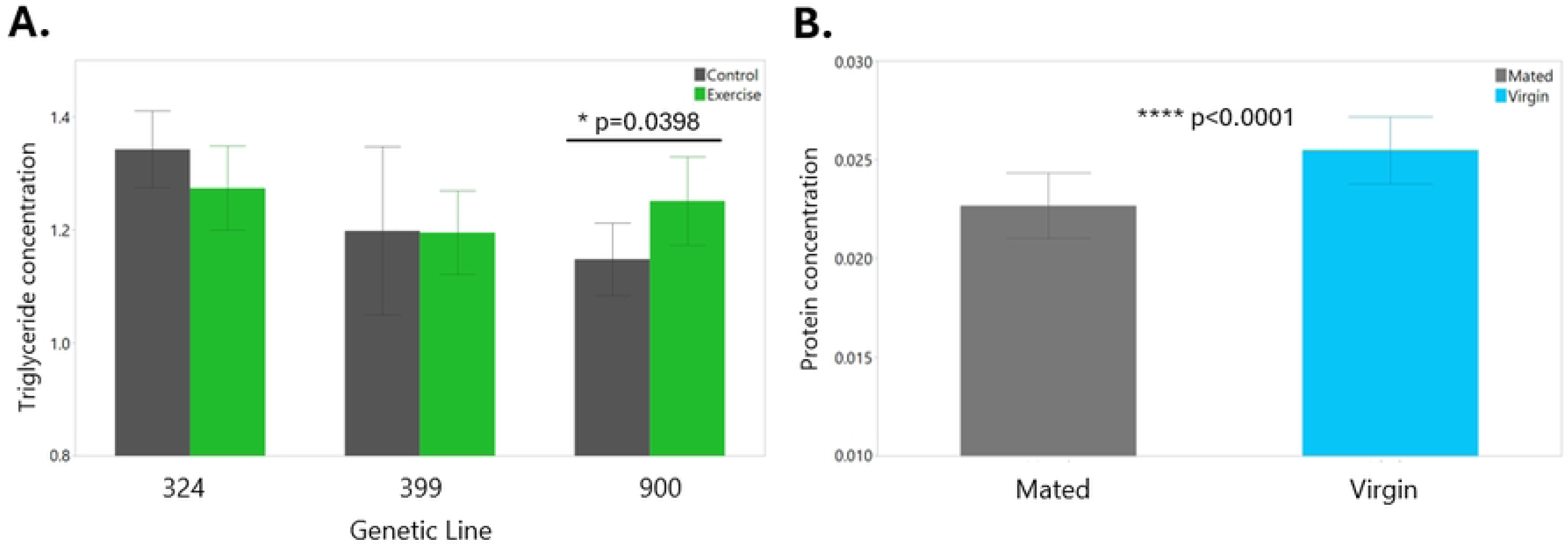
Genetic line and exercise influence triglyceride concentrations, while mating status influences protein concentrations. (A) Normalized triglyceride concentration results for each exercise treatment by genetic line. Error bars are one standard deviation. (B) Protein concentration results for mated (gray) and virgin (light blue). Error bars are one standard deviation.

Finally, we analyzed the expression of four genes to gain a better understanding of some of the molecular differences between Control/Exercised and mated/virgin flies. We chose to measure the expression of *Tdc1* and *Tbh*, which encode the enzymes necessary for OA synthesis. We also chose *Dilp2* (one of the main insulin-like peptides in *Drosophila*) and *AkhR* (the receptor for adipokinetic hormone (Akh)), which are both known to be regulated by OA signaling [28]. Interestingly, we observed no differences in gene expression between the Control and Exercised flies, even though another study found that *Tdc1* and *AkhR* are upregulated in larvae post-exercise [38]. However, we did find that virgin female flies had increased expression of *Dilp2*, *Tbh*, and *AkhR* (Fig 6).

**Fig 6.**
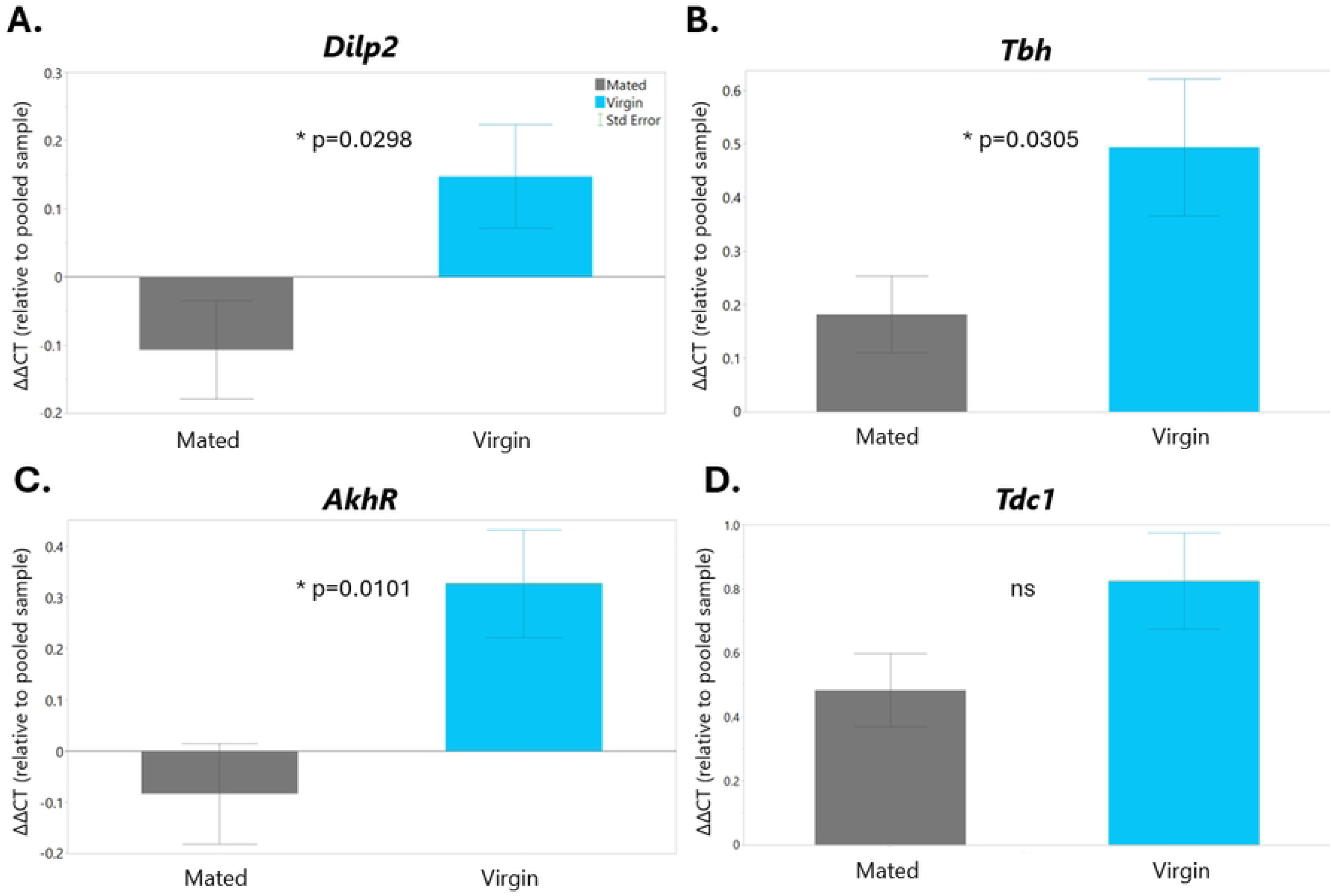
Virgin female flies have upregulated *Dilp2*, *Tbh*, and *AkhR* expression levels. (A) Relative gene expression levels for *Dilp2*. Mated females are shown in gray and virgin females are shown in light blue for all graphs. (B) Relative gene expression levels for *Tbh*. (C) Relative gene expression levels for *AkhR*. (D) Relative gene expression levels for *Tdc1*. All error bars are standard error.

## Discussion

Exercise is an example of hormetic stress, which is defined as the phenomenon where repeated exposure to a moderate stressor causes physiological adaptations that allow the organism to better respond to other stressors [39]. Subsequently, it makes sense that systems such as the Fight-or-Flight stress response are critical for exercise adaptations. In *Drosophila*, the main hormone/neurotransmitter that regulates the Fight-or-Flight stress response is octopamine, which has previously been implicated in *Drosophila* exercise adaptations [24,40]. Human studies have also confirmed the importance of norepinephrine/epinephrine in exercise adaptations. At the onset of exercise, their circulating levels increase to elevate heart rate and blood pressure, as well as to mobilize glucose and free fatty acids to be used as fuel [41]. Furthermore, patients taking beta blocker medications (which block epinephrine/norepinephrine from binding to beta adrenergic receptors) have reduced exercise performance, underscoring the importance of these hormones in exercise response [42]. However, human studies have had conflicting results regarding how sex can influence epinephrine/norepinephrine levels and exercise performance [41].

The goal of our research study was to extend previous results in the *Drosophila* exercise field to understand why we observe such a large difference in how male and female flies respond to exercise. Contrary to previous studies, our results from Experiment 1 suggest that an exercise program of one hour of exercise per day for five days can increase mated female flies’ climbing performance but has no effect on male flies. When we increased the exercise duration to two hours in Experiment 2, our findings showed that Exercised mated females had decreased climbing performance compared to Control. It could be possible that shorter, less intense exercise training could be beneficial for mated females, but increasing the duration can have negative effects. We also performed the climbing assay at different time points post-exercise for the two experiments (two days post-exercise treatment for Experiment 1, and nine days post-exercise treatment for Experiment 2), which could also be a confounding factor. Overall, our results suggest that exercise response in *Drosophila* is highly context specific, with duration of exercise, exercise device, genotype, and sex all playing a role.

We hypothesized that a female’s mating status would influence how they respond to the stress of exercise, and that exercise would affect fertility. Our data showed that genotype and mating status had statistically significant effects on fitness traits such as climbing performance, lifespan, and fertility. Virgin female flies were able to climb higher than mated female flies; however, they had a significantly shorter lifespan. Contrary to our present findings, several previous studies have found that mating is associated with a decreased lifespan for females referred to as the survival cost of reproduction [43,44]. Mating, especially with multiple males, can cause physical damage to females which increases their mortality [45]. Moreover, female flies selected for longevity have increased fertility later in life [46]. However, other studies have found that this survival cost of reproduction is not always repeatable and can vary based on genotype and experimental conditions [47]. We conclude that these reasons could be why we observe an opposite pattern when comparing the lifespan of virgin and mated females. Additionally, we found that mated female flies who were subjected to exercise had fewer offspring compared to Control, suggesting that exercise exerts harmful levels of stress on these flies. We had hypothesized that exercise was causing them to use up their energetic stores with not enough left over to produce offspring, but we found that exercise had no effect on triglyceride or protein levels after nine days of rest.

We also performed qPCR to gain an understanding of how exercise and/or mating status might affect flies on a molecular level and found that virgin female flies had upregulated expression of *Dilp2*, *Tbh*, and *AkhR*. These results suggest that virgin flies have increased levels of both insulin signaling and octopamine signaling, which is interesting as these have opposing effects on metabolism. *Dilp2* acts to reduce glucose levels in the hemolymph by increasing glucose uptake in cells, while OA acts to break down energy stores to increase glucose and free fatty acids in the hemolymph [28,48]. In addition, a previous study had found that increased OA levels are associated with decreased *Dilp2* release [27]. Increased expression of *Tbh*, which encodes the enzyme necessary for OA synthesis, likely indicates that virgin flies are mounting a higher stress response than mated female flies. Additionally, increased insulin signaling is associated with decreased lifespan [49], which is consistent with our findings that virgin female flies have shorter lifespans than mated female flies. Overall, our results suggest that virgin females have increased OA signaling, and thus may be mounting an increased stress response, possibly leading to the activation of Akh signaling to break down stored fats into energy. Further investigation involving more direct measures of these signaling pathways is required to draw more substantial conclusions.

We conclude that mating status dictates how female flies respond to stress. It is difficult to say whether mated or virgin females are more or less vulnerable to the stress of exercise, as they seem to respond differently in all the phenotypes we measured. Female flies under stress likely experience an evolutionary trade-off where it is advantageous for virgins to expend more energy being active, while mated females conserve their energy and are able to survive longer so that they can reproduce.

## Conclusion

This study sought to explore the interactions between mating status, physical exercise, and octopamine in *Drosophila*. Our findings confirm that sex and genotype strongly influence flies’ ability to adapt to exercise, but we found that OA supplementation had little effect. We also show that virgin females have greater climbing abilities than mated females post-exercise, although this seems to come at a cost to their survival. Exercise training appears to have a negative effect on fertility, with effects lasting a week after the cessation of exercise. However, there were no major effects of exercise treatment or mating status on triglyceride levels. Exercise training also had no significant effect on expression of the genes we examined, but virgin females had increased transcript levels of *Dilp2*, *Tbh*, and *AkhR* compared to mated females. Taken together, these results suggest that mating status is an important indicator of how female flies will respond to exercise, and that there are likely evolutionary tradeoffs occurring in these flies. We believe this research will lay the groundwork for future studies to further explore sexual dimorphism and the effects of mating on exercise response.

## Acknowledgements

We would like to thank Tolulope R. Kolapo, Ryan L. Earley, and Stanislava Chtarbanova for their helpful advice on this study.

